# Stereotypic expansion of T_regulatory_ and Th_17_ cells during infancy is disrupted by HIV exposure and gut epithelial damage

**DOI:** 10.1101/2021.05.03.442468

**Authors:** Sonwabile Dzanibe, Katie Lennard, Agano Kiravu, Melanie S.S. Seabrook, Berenice Alinde, Susan P. Holmes, Catherine A. Blish, Heather B. Jaspan, Clive M. Gray

## Abstract

Few studies have investigated immune cell ontogeny throughout the neonatal and early paediatric period, where there is often increased vulnerability to infections. Here, we evaluated the dynamics of two critical T cell populations, regulatory (Treg) cells and Th17 cells, over the first 36 weeks of life. Firstly, we observed distinct CD4^+^ T cells phenotypes between cord blood and peripheral blood, collected within 12 hours of birth, showing that cord blood is not a surrogate for newborn blood. Secondly, both Treg and Th17 cells expanded in a synchronous fashion over 36 weeks of life. However, comparing infants exposed to HIV *in utero*, but remaining uninfected (iHEU), with HIV-unexposed uninfected control infants (iHUU), there was a lower frequency of peripheral blood Treg cells at birth, resulting in a delayed expansion, and then declining again at 36 weeks. Focusing on birth events, we found that Treg cells co-expressing CCR4 and α4β7 inversely correlated with plasma concentrations of CCL17 (the ligand for CCR4) and intestinal fatty acid binding protein (iFABP), IL-7 and CCL20. This was in contrast to Th17 cells, which showed a positive association with these plasma analytes. Thus, despite the stereotypic expansion of both cell subsets over the first few months of life, there was a disruption in the balance of Th17 to Treg cells at birth likely being a result of gut damage and homing of newborn Treg cells from the blood circulation to the gut.

**Key points:**

1. Phenotypic differences between cord and birth peripheral blood CD4 cells.
2. Synchronous increase of Th17-Treg cells is disrupted by HIV/ART exposure.
3. Intrauterine HIV exposure was associated with epithelial gut damage.

## Introduction

Early infancy has the highest risk of mortality in children under 5 years, and with over 45% of these deaths occurring in the first month of life (1). Neonates are more likely to succumb to infections compared to adults, largely due to their cellular immune system being less effective at protecting against invading pathogens (2). Although the foetal immune system has matured by 15 weeks’ gestation, foetal immunity exhibits heightened tolerogenic activity, Th2/Th17 polarisation bias, and lacks antigen experience which can result in blunted cellular immunity to pathogenic insults in early perinatal life (3–6). Following delivery, the neonatal immune system has to abruptly undergo adaptations to cope with extrauterine pathogens whilst developing tolerance to oral antigens and commensal microbes. Since postnatal immune interactions determine risk to subsequent infections and immune-related disease (7, 8), understanding longitudinal ontogeny and factors that may disrupt development of the immune system are of critical importance in designing novel treatments and vaccine strategies to combat the high burden of disease in neonates.

CD4^+^ helper T cells influence immunological responses to antigens either by promoting inflammation or establishing tolerance. T cell adaptive immunity among infants tends to exhibit weak Th1 polarization capacity (6) and thus the homeostatic balance and interplay between Th17 and Treg cells during an inflammatory response is key to controlling infection while limiting immunopathology (9). The ontogeny of these immune cells early in life is therefore crucial in establishing balanced immune responses towards pathogens, commensals and autoantigens. There are, however, limited studies reporting on the longitudinal development of Th17 and Treg cells, hence impeding potential corrective intervention strategies to ensure optimal protective immunity so being able to reduce the burden of infectious disease early in life.

There is a growing population of infants born to women living with HIV globally (10), and although prenatal antiretroviral treatment (ART) has reduced mother to child HIV transmission, HIV-exposed uninfected infants (iHEU) experience 60-70% higher mortality risk compared to their HIV-unexposed uninfected (iHUU) counterparts (11). Infectious diseases are the most likely cause of iHEU mortality, owing to higher rates of gastrointestinal and respiratory tract infections compared to iHUU (12–14). It is possible that maternal HIV infection can influence the Th17 and Treg immune balance in these children at birth and upset the trajectory of inflammatory versus tolerogenic immune cell ontogeny during early life. Thus, understanding the balance between immune inflammation and regulation in iHEU may also shed light on immune disturbances during infancy in general, and would also lay a foundation for providing more informed health-improving treatments for these vulnerable children.

Therefore, we aimed to define the ontogeny and relationship between Th17 and Treg cells in the first 9 months of life and examine how HIV/ARV exposure could impact the trajectory of these immune subsets. Here we show that the most dramatic change in the immune system occurs during the early period postpartum. Our data reveal that HIV/ARV exposure perturbs immune homeostatic and/or regulatory balance early in life, likely because of impaired epithelial gut integrity in iHEU compared to iHUU.

## Materials and Methods

### Study participants

We conducted a longitudinal study on the effects of *in utero* HIV exposure on the phenotypic development of CD4^+^ T lymphocytes during early infancy. Women were recruited to participate in the study within few hours following delivery as described (15). Women aged ≥18 years, who experienced no complications during pregnancy and provided signed informed consent for themselves and their respective infants were enrolled into the study approved by human research ethics committee of University of Cape Town (FWA1637; IRB0001938). Infants with gestational age <36 weeks and birth weight <2.4 kg were excluded from study analysis. Peripheral blood was collected from the infants at birth (<12 h), 7, 15 and 36 weeks of infant age. Among infants born to mothers living with HIV, absence of HIV transmission was confirmed by performing HIV DNA PCR test at 6 weeks of infant age. All infants received childhood vaccines according to the South African extended program of immunization schedule. A total of 36 infants; 20 iHEU and 16 iHUU were included in this analysis and followed over 36 weeks of life. To reduce potential confounding factors, samples were selected that showed no demographic differences between the two groups of infants (Table S1). All mothers elected to exclusively breastfeed at birth, although by 7 weeks only 52.8% were determined to be still exclusively breastfeeding (Table S1). All mothers who tested positive for HIV received combined (c)ART as recommended by WHO guidelines (16). Of the 20 iHEU, 10 (50%) were born to HIV-infected women who initiated cART during pregnancy at a median of 24.9 weeks gestation. We termed these infants iHEU-i and would likely have had more potential HIV exposure in the first two trimesters than the remaining women, who initiated cART prior to conception. The infants born to these women were termed iHEU-s (Table S1).

### Whole blood collection

Blood was collected from the infants directly into sodium heparin tubes and transported to the laboratory for processing within 6h. BD FACS Lysing Solution (BD Biosciences, CA USA) was used for lysing of red blood cells and the fixation of the remaining peripheral blood mononuclear cells (PBMC) according to manufacturer’s instructions. Fixed whole blood cells were stored at −80°C for ~24 h in foetal calf serum with 10% DMSO and then cryopreserved at −180°C until assayed using flow cytometry. Plasma samples were stored at −80°C until used for serological analysis in this study.

### Flow cytometry analysis

Immunophenotyping of CD4^+^ Th17 and Treg cells in infants was performed by staining thawed cells using a multi-colour antibody panel (Table S2). Lymphocytes were characterized by surface staining with antibodies specific to CD3, CD4 and CD8. To delineate Treg cells from the lymphocyte population, antibodies specific to surface makers CD127, CD25, CD39 and TIGIT together with transcription factor FoxP3 were used. Supplementary Figure S1 shows the gating strategy that was used to define Treg cells, showing that the highest expression of Treg markers (FoxP3, CD39 and TIGIT) was found in the CD4^+^CD25^hi^CD127^-^ population compared to either CD4^+^CD25^lo^CD127^-^ or CD4^+^CD25^lo^CD127^+^ populations. Surface staining of CCR6, CCR4 and CD161 was used to define Th17 cells. Using the manual gating approach, we therefore defined Treg cells as: CD4^+^CD25^+^CD127^-^FoxP3^+^ and Th17 cells as CD4^+^CCR6^+^CCR4^+^CD161^+^. Gut homing cells were defined using anti-α4β7-surface staining. Immune cells migrating to the lymph nodes were characterized using CCR7 specific antibodies. Data acquisition was performed using BD LSR II and analysed with FlowJo software (version 10.5.3, Tree Star Inc., CA).

### Quantification of intestinal fatty acid binding proteins using ELISA

Infant plasma samples collected at birth, 7, 15 and 36 weeks of age were used to measure the concentration of intestinal fatty-acid binding protein (iFABP) using a commercial ELISA kit (Elabscience). Test samples were diluted 1/400 in assay diluent and measured in duplicates according to manufacturer’s instructions. An in-house high titre control plasma sample was included to determine intra-assay variation which had a coefficient of variation = 5.7%, indicating good intra-assay comparisons.

### Quantification of chemokine and cytokine concentrations using multiplex immunoassay

A 10-plex immunoassay was designed to measure plasma concentrations of chemokines and cytokines in infant plasma samples collected at birth and 36 weeks of age. Using the multiplex immunoassay kit (R&D systems, Minneapolis, MN), we measured CCL17, CCL22, CXCL10 (IP10), CCL20 (MIP3A), CCL25, IL-6, IL-7, IL-17A, IL-10 and IL-27 concentrations according to manufacturer’s instructions. Test samples were diluted 3:1 with assay diluent and signal intensity measured using Biorad-200 Luminex platform.

### Statistical analysis

R software (version 3.5 R Core Team, Vienna, Austria) was used to test for statistical differences. The cell populations described in the study are reported as median percentages. iFABP concentrations were log10 transformed for statistical comparisons. Nominal variables were compared using χ^2^ or Fisher exact test. Differences in cell frequencies between the two groups of infants were compared using the Mann-Whitney *U* test. Friedman’s test was used to compare differences within a group over time. Chemokine and cytokines included in a multiplex immunoassay were compared using Mann-Whitney test and Bonferroni correction used to adjust for multiple comparisons.

Coordinates for multidimensional scaling of markers expressed on CD4^+^ cells were calculated using metaMDS function and PERMANOVA statistical differences performed using the adonis function from the vegan R package. Generalized linear mixed model (GLMM) analysis was performed using an open-source R package CytoGLMM as described in (17). Unsupervised cell population identification was performed using self-organizing map and hierarchal clustering as implemented in the FlowSOM and metaclustering R packages (18). This approach not only allowed us to identify novel populations of Treg and Th17 clusters of cells but also avoid the bias of cell identity due to manual gating of known populations. The mixOmics R package was used to integrate the identified CD4^+^ cell clusters with the measured plasma analytes. RV coefficients were computed, and the latent variables determined by sparse partial least squares (sPLS). To determine the strongest predictors of HIV exposure at birth and week 36 we performed sPLS discriminant analysis (sPLS-DA) and computed the loading plot and ROC plot of component 1 of all variables in our data set (19). p-value < 0.05 was considered statistically significant.

## Results

### Stereotypic expansion of Th17 and Treg cells from birth to 36 weeks

There is a paucity of data describing the ontogeny of Th17 and Treg cells throughout the first few months of life, with most studies comparing cord blood with infant and adult peripheral blood (20–22). To describe the ontological changes occurring during infancy, we compared matched infant peripheral blood samples collected at birth, 7, 15 and 36 weeks of age from 16 healthy, term, breastfed infants. A total of sixty-four matching infant blood samples were analyzed by flow cytometry and used manual gating to determine the proportion of CD4^+^CCR6^+^CCR4^+^CD161^+^Th17 and CD4^+^CD25^hi^CD127^-^ FoxP3^+^ Treg cells and reported as the percentage of the parent CD4^+^ population (Figure S1). The proportions of Th17 cells were lowest at birth and gradually increased with age, peaking at 15 weeks (Figure 1A). Treg cells shared a similar stereotypic change as Th17 cells, being lowest at birth and increasing with age, although by 7 weeks the levels stabilized, and a marginal decline was observed by 36 weeks (Figure 1B). The similar stereotypic changes of Th17 and Treg cells in the first 36 weeks of life was reflected in the significant correlation found between these two populations (Figure 1C). Knowing that Treg and Th17 cells exhibit high plasticity and in specific conditions have been shown to undergo trans-differentiation between the two CD4 subsets (23, 24), we examined whether the co-linearity of these cells was due to marker co-expression. Using uniform manifold approximation and projection (UMAP) analysis, Treg and Th17 markers were expressed on distinct subsets of cells (Figure 1D). Our data reveal that circulating Treg and Th17 cells show similar stereotypic changes during infancy and that the parallel changes of Treg and Th17 cells are not due to a co-lineage of cells but are distinct populations.

**Figure 1:**
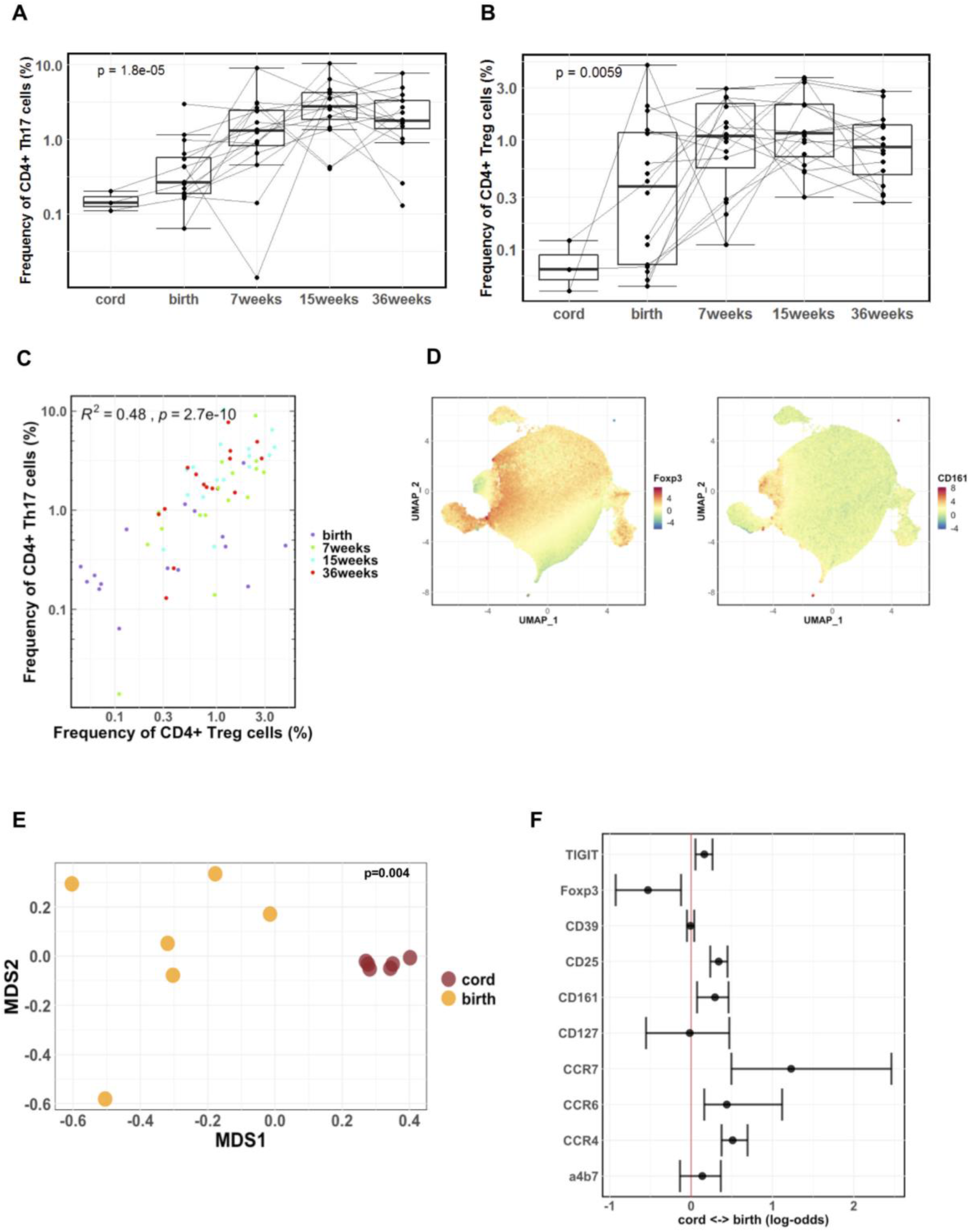
Stereotypic expansion of Th17 and Treg cells from birth to 36 weeks. A & B) Changes in the frequency of Th17 and Treg cells from birth to 36 weeks of age, p value show Friedman’s test across time points. C) Spearman rank correlation between Th17 and Treg cells across birth, 7, 15 and 36 weeks. D) UMAP analysis displaying Foxp3 and CD161 expressing CD4^+^ T cells distinguishing the lineage of Treg cells and Th17 cells. E) Multidimensional scaling (MDS) showing dissimilarity of CD4^+^ cells from cord and peripheral blood at birth measured by PERMANOVA. F) Generalized linear mixed model comparing CD4 marker expression between cord blood and peripheral blood at birth. The red horizontal line denotes zero difference and solid symbols represent median differences ± interquartile ranges.

### Cord blood and birth peripheral blood CD4 T cell populations are distinct

Cord blood is frequently used as a proxy for infant blood in early life studies of the immune system for logistical reasons. Since the proportions of Th17 and Treg cells were lower in cord blood compared to birth blood (Figure 1A & B), we determined the similarity between cord blood and infant peripheral blood collected at birth. We used a polychromatic flow cytometry panel (Table S1) to analyze CD4^+^ phenotypes, in a limited set of 6 matching cord blood samples and peripheral blood collected within the first 12 hours of life. First, we performed multidimensional scaling (MDS) on the fluorescent intensities of the cell markers. CD4^+^ cells from cord blood uniquely clustered distinctly from birth blood samples (Figure 1E). To determine which cell markers distinguished cord and birth blood CD4 cells, we used generalised linear mixed model (GLMM) regression that calculates log-odds expression to identify markers predictive of either blood type (17). There was a significant increased odds of birth peripheral blood CD4^+^ cells expressing CCR7, CCR4, CCR6 and CD25 compared to cord blood (Figure 1F). Conversely, cord blood CD4^+^ cells had increased odds of expressing FoxP3 compared to birth blood (Figure 1F). These data show that there are phenotypic differences of CD4^+^ T cells between cord blood and birth blood and, despite the limited sample size, suggests that cord blood should not be equated with birth blood and that rapid changes in circulating T cells occur within hours of delivery.

### Identity of heterogenous CD4 clusters during infancy

Treg and Th17 cells are not necessarily a defined static population and varying phenotypes have been described that play specific immunological roles (25, 26). To further define the heterogeneity of inflammatory versus regulatory CD4 cells during infancy in an unbiased manner; we used unsupervised Flow Self-Organizing Maps (FlowSOM) and hierarchical clustering to identify unique CD4 clusters of cells. This is an unbiased and deeper phenotyping approach, based on the expression of all markers included in the flow panel (Table S2), and can identify more nuanced and novel populations that exist within the conventional manual gates, and would hence be missed. Using this approach, we identified 9 CD4^+^ T cell clusters. Figures 2A & B show delineation of the CD4^+^ populations by phenotype. Cell cluster relatedness is displayed using hierarchal clustering dendrograms and mapping the cell clusters on UMAP (Figure 2C). We identified two clusters of Treg cells, a small population of CCR4^+^ Tregs (cluster 3, Figure 2B) and relatively larger population of α4β7^+^ Tregs (cluster 5, Figures 2A & B).

**Figure 2:**
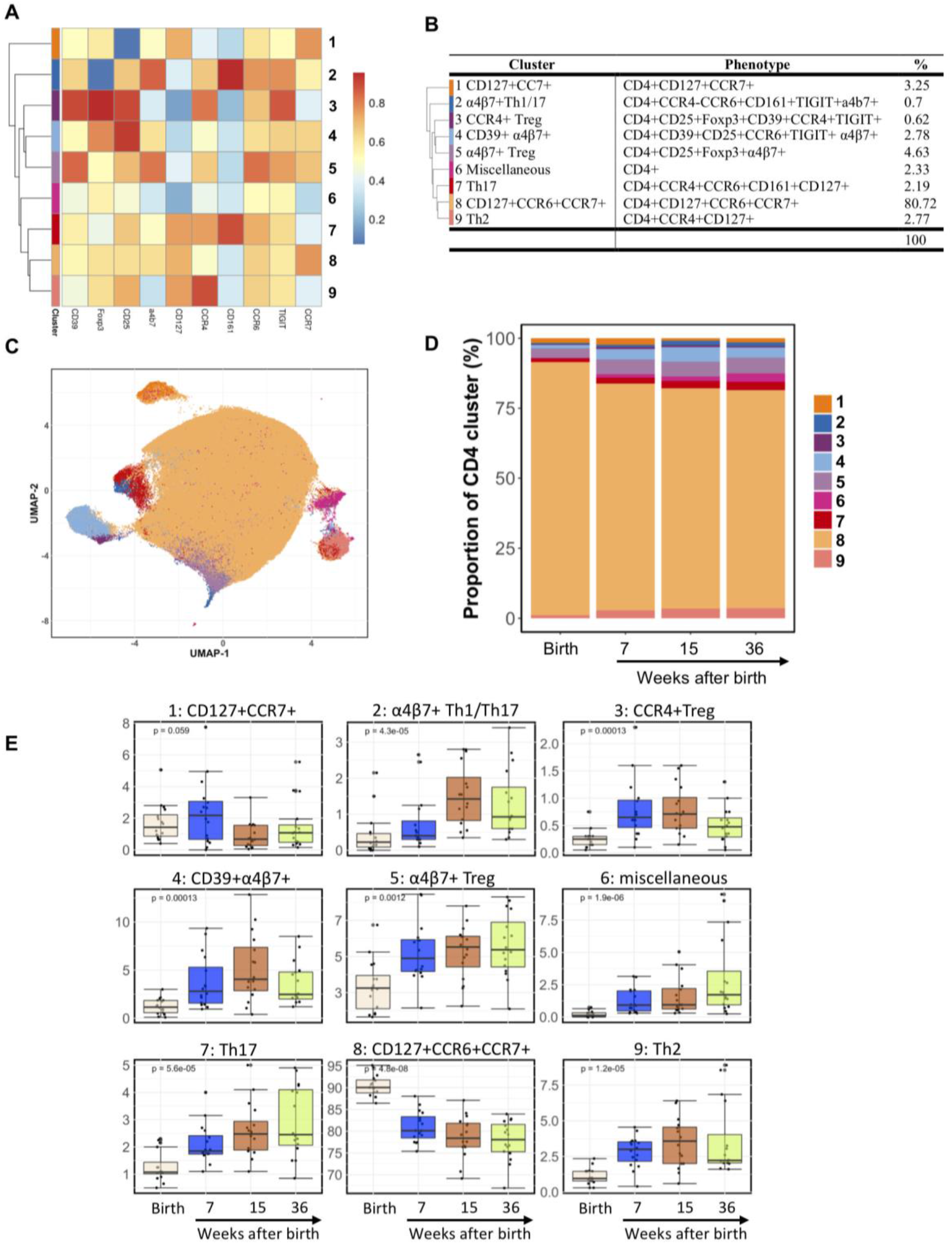
Identity of heterogenous CD4 T cell clusters during infancy. A) Hierarchical clustering showing 9 CD4^+^ T cell clusters identified using FlowSOM. B) Listing of the 9 clusters by phenotype, showing the proportions of each cluster identified using FlowSOM. C) Dimensional reduction projecting each of the 9 clusters in relation to each other on the UMAP. D) Relative abundance of CD4^+^ T cell clusters at birth, 7, 15 and 36 weeks. E) Box and whisker plots showing individual changes over infant age for each CD4^+^ T cell cluster, p value show Friedman’s test across time points.

The CCR4^+^ Treg cluster 3 co-expressed ectonucleotidase CD39, known to disrupt metabolic activity of activated and proliferating cells (27), and the inhibitory molecule, TIGIT, that prevents cognate interaction between antigen presenting cells and T helper cells (28), suggesting that the CCR4^+^ Treg cluster 3 was a highly suppressive Treg population (Figure 2A & B). The 7-fold larger population of α4β7^+^ Tregs (cluster 5) co-expressed CCR7^+^, characteristic of gut trafficking lymphocytes (Figure 2A & B). Both clusters 3 and 5 were distinct from the 7 other CD4+ population clusters but did show a close relationship on the UMAP (Figure 2C). Figures 2D and E show the trajectory and ontology of the 9 CD4 populations, where Figure 2D shows the clusters by order of proportions and Figure 2E shows the detailed change from birth. Except for clusters 1 and 8, cell populations increased following birth and remained relatively stable at later time points although by 36 weeks, the proportion of CCR4^+^ Treg cluster 3 significantly decreased (Figure 2E). We also identified a Th17 cluster (7, CCR6^+^CCR4^+^CD161^+^CD127^+^) that increased with advancing infant age and cluster 2, denoted as Th1/17 α4β7^+^ (CCR6^+^CCR4^-^CD161^+^) which increased with infant age and peaked at 15 weeks (Figure 2D & E).

The trajectory of these clusters from birth is shown in Figure 2E, revealing that several CD4^+^ populations contracted and expanded following birth. A naïve-like CD4+ T cell cluster 8 (CD127^+^CCR6^+^CCR7^+^), accounting for >80% of the CD4 cells at birth, significantly declined over time, presumably as the other antigen-experienced cells expanded (Figure 2E). Only one CD4^+^ cluster (1, CD127^+^ CCR7^+^) remained relatively stable from birth up to 36 weeks (Figure 2E). The other Treg and Th17 clusters all expanded after birth, including a likely Treg suppressive gut-homing CD39+ α4β7^+^ population: 4, CD39^+^CD25^+^CCR6^+^TIGIT^+^α4β7^+^; a non-migratory CD39+TIGIT-expressing Treg population: 3, CD25+FoxP3+CD39+CCR4+TIGIT+; a potential less-suppressive Treg migratory population: 5, CD25+FoxP3+α4β7^+^. Non-Treg population expansions included a Th2 cluster (9, CCR6^-^CCR4^+^), and a miscellaneous cluster 6 that did not express any of the markers analysed (Figure 2D and E). Collectively, these findings show an expansion of differentiated CD4^+^ T cell subsets with age compared to the decreasing frequency of naïve cells after delivery (20). The most dramatic changes were observed to occur in the first 7 weeks of life, a period of immune plasticity as the adaptive immune system is likely shaped by environmental antigens postpartum (7, 8).

### Disrupted Th17-Treg ratio in iHEU

We next parsed out differences between CD4^+^ clusters in iHEU from iHUU, as it is known that there is a more activated threshold of pro-inflammatory cytokine production from innate immune cells (29) and skewed adaptive immunity towards activated and exhausted cells in iHEU (30–32). To assess whether these known alterations might be related to the Th17-Treg ratio, we compared the phenotype of CD4^+^ cells of the 16 healthy infants (iHUU) with that of 20 iHEU. MDS of marker expression revealed distinct clustering of CD4^+^ T cell marker expression between iHEU vs iHUU, regardless of age (Figure 3A). The observed differences were driven by significantly higher odds of Foxp3 and CD161 expression in iHUU compared to iHEU, which persisted until 36 weeks of age (Figure 3B). By 36 weeks, log-odds expression of α4β7 was also dependent on HIV exposure, being higher in iHEU compared to iHUU (Figure 3B). These findings show that CD4^+^ T cells of iHEU can be distinguished from that of iHUU primarily by higher expression of Foxp3 and CD161, which are Treg and Th17 discriminatory markers, respectively.

**Figure 3:**
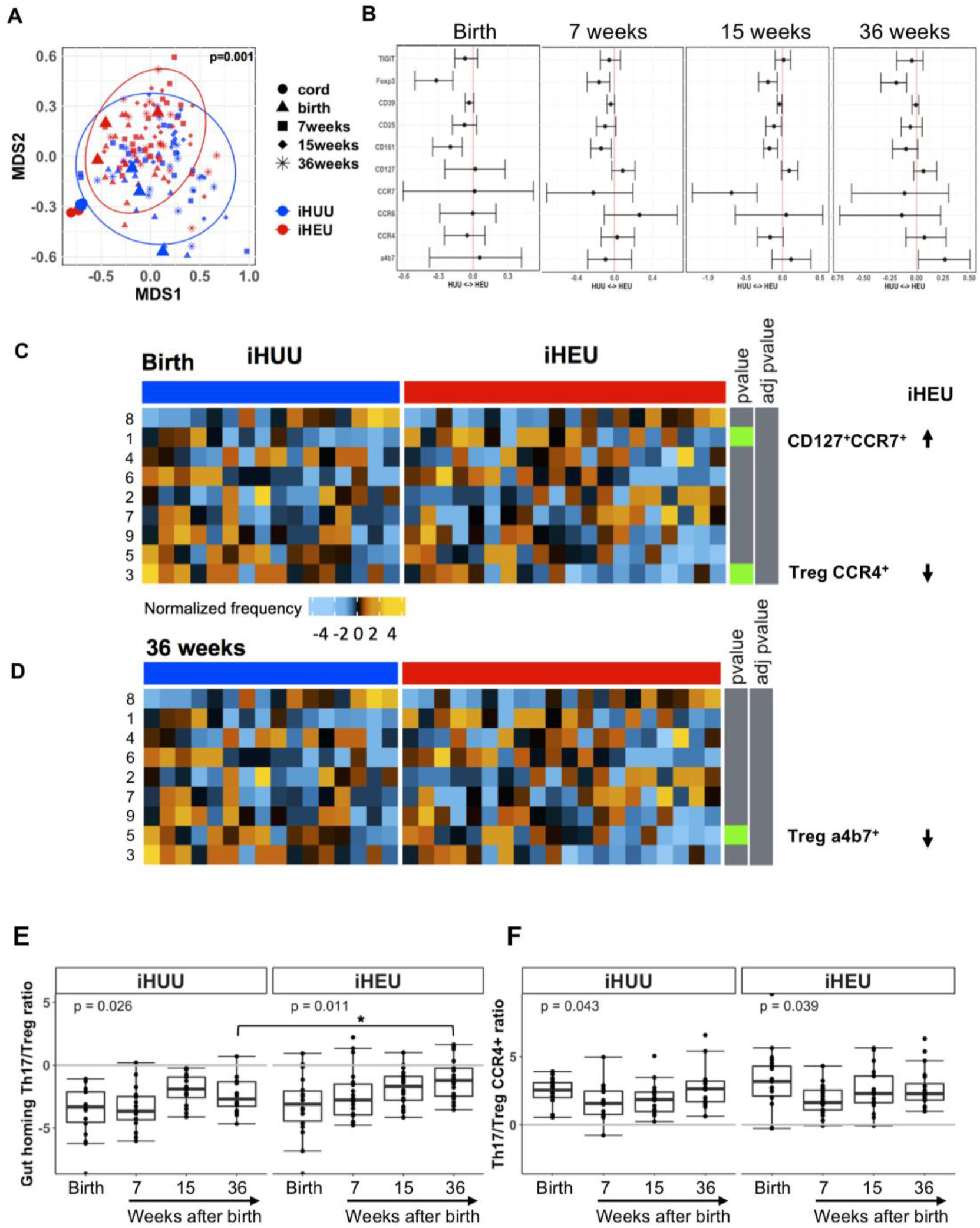
The impact of HIV exposure on proportions of CD4+ T cell clusters and the Th17 and Treg balance. A) Multidimensional scaling (MDS) showing dissimilarity of CD4^+^, measured by PERMANOVA, between HIV-exposed uninfected infants (iHEU) and HIV-unexposed uninfected infants (iHUU). B) Generalized linear mixed model (GLMM) comparing CD4 marker expression between iHEU and iHUU. C & D) Differential abundance of CD4 cell clusters in iHUU and iHEU at birth and 36 weeks compared using GLMM and false discovery rates (FDR) used to adjust for multiple comparisons. Green squares indicate a significant p-value. Arrows indicate whether the population cluster was increased or decreased in iHEU. E&F) Log2 Th17/Treg ratio, Friedman’s test used to show changes within a group and *p<0.05 show differences between iHEU and iHUU compared using Mann-Whitney *U* test.

We next performed differential abundance testing to assess whether HIV exposure alters the proportion of CD4^+^ cell clusters defined in Figure 2A. When examining the proportions of Treg and Th17 CD4^+^ cell clusters between iHUU and iHEU at birth (Figure 3C), the minor, more highly suppressive CCR4^+^ Treg cluster 3 was lower in iHEU compared to iHUU (p=0.015), albeit not statistically significant after adjusting for multiple comparisons (Figure 3C & S2A). Conversely, the CD4^+^CD127^+^CCR7^+^ T cell cluster 1 was significantly higher in iHEU (p=0.027), although losing significance after adjusting for multiple comparisons (Figure 3C). At 36 weeks (Figure 3D), the α4β7^+^ Treg cluster 5 was significantly lower in iHEU (p=0.029), also losing significance after adjustment. Our manual gating approach revealed a similar pattern, where several of the CD4^+^ T cell clusters identified by SOMs would fall within the Treg gate (all CD127^-^ CD25^+^FoxP3^+^, including CCR4^+^ and α4β7^+^ Treg) or the Th17 gate (all CD161^+^CCR4^+^CCR6^+^, including CD127^+^CCR7^+^). For example, there was a significantly lower proportion of Treg cells at birth (p=0.017) and at week 36 (p=0.012; Figure S3B). These changes translated to an offset of the Th17/Treg balance in iHEU, where the ratio increased with infant age (p=0.015; Figure S4) compared to a relatively stable Th17/Treg ratio in iHUU (p=0.21; Figure S4). Similarly, the ratio of gut homing (α4β7^+^) Th17/Treg cells steadily increased with infant age in iHEU (Figure 3E), resulting in a higher Th17/Treg ratio in iHEU by 36 weeks (Figure 3E) and which was significantly higher in iHEU versus iHUU (Figure 3E). This increase in the Th17/Treg ratio was not observed to the same degree when gating on CCR4+ Th17 and Treg cells (Figure 3F), suggesting that the changes over time in iHEU was driven by α4β7^+^ expressing Treg cells. Even though blood circulating Th17 and Treg cells show similar stereotypic changes during infancy, the observed Th17/Treg imbalance in iHEU are due to depressed frequencies of Treg cells.

When we stratified iHEU based on the timing of maternal cART initiation, as a proxy for potential *in utero* HIV exposure, there was no statistically significant difference between Treg and Th17 CD4+ cell clusters or phenotypes between iHEU-s and iHEU-i using either our SOM or manual gating approach (Figures S2B and S3C & D).

### Treg and Th17 proportions associate with iFABP and gut-tropic chemokines and cytokines

Previous reports of intestinal damage leading to inflammation in iHEU (33) led us to investigate the relationship between our different phenotypic clusters of cells with intestinal fatty-acid binding protein (iFABP) plasma concentrations. iFABP concentrations are elevated when there is impaired gut epithelial integrity leading to microbial translocation and gut inflammation (34). Indeed, infant plasma concentrations of iFABP at birth were significantly higher in iHEU compared to iHUU (Figure 4A). By 36 weeks of life, no differences in iFABP concentrations were observed between iHEU and iHUU (Figure 4A), nor at weeks 7 and 15 (Figure S5A). We also stratified by potential HIV exposure, as measured by initiation of ART and found there to be no difference in plasma iFABP concentrations between iHEU-s and iHEU-I (Figure S5B). Of the 10 plasma concentrations of inflammatory and regulatory chemokines and cytokines at birth and 36 weeks (Figure S5C), CCL17 was significantly higher in iHEU compared to iHUU at birth (Figure 4B). The regulatory cytokine, IL-27, involved in epithelial restitution (35) was observed to be higher in iHEU, albeit no longer significant after adjusting for multiple comparisons (Figure 4B) As with iFABP, there was no difference between the 10 analytes when parsed between iHEU-s and iHEU-i (Figure S5D).

**Figure 4:**
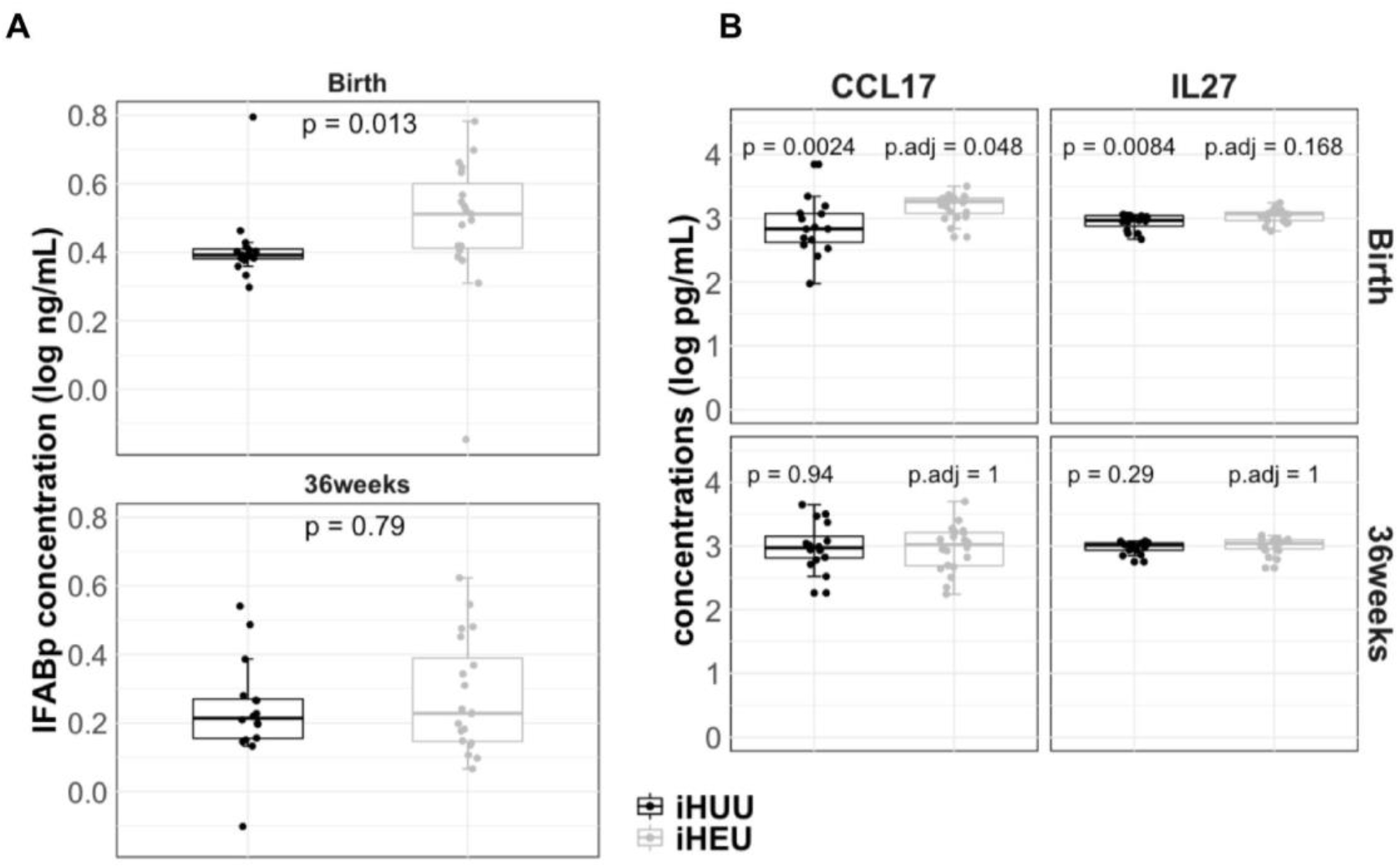
Plasma concentrations of intestinal fatty acid binding protein (iFABP), CCL17 and IL-27 at birth and 36 weeks in iHUU and iHEU. A) Comparing iFABP plasma concentrations between HIV-unexposed uninfected infants (iHUU) and HIV-exposed uninfected infants (iHEU) using the Mann-Whitney *U* test. B) Comparing CCL17 and IL-27 plasma concentrations in iHUU and iHEU at birth and 36 weeks using the Mann-Whitney *U* test and Bonferroni adjusted p values.

To integrate the multimodal cellular and protein data, we used generalized canonical regression to calculate RV coefficients of the latent variables determined by sparse partial least squares (sPLS). Using this approach, we assembled cluster image maps showing the association between CD4^+^ clusters and plasma analytes at birth and 36 weeks (Figure 5A-D). At birth (Figure 5A) the Treg clusters shared a negative association with CCL17, iFABP, IL-7 and CCL20, which was in stark contrast to Th17 and CD127^+^CCR6^+^CCR7^+^ clusters, showing a positive association (Figure 5A). More precisely, the frequencies of Treg clusters (CCR4^+^ Treg and α4β7^+^ Treg) were inversely correlated with iFABP (RV= −0.36 and −0.35 respectively), IL7 (RV=-0.37 and −0.37 respectively), CCL17 (RV= −0.36 and −0.36 respectively) and CCL20 (RV= −0.35 and −0.34 respectively) concentrations, while the frequency of Th17 cluster was positively correlated to iFABP (RV = 0.25), IL7 (RV= 0.26), CCL17 (RV= 0.25) and CCL20 (RV=0.24) (Figure 5A). Furthermore, using sPLS discriminant analysis (PLS-DA), we were able to determine the weighted score of each variable to predict HIV exposure: CCL17, IL27, CCR4^+^ Tregs, CD127^+^CCR7^+^ T cells and iFABP were the strongest predictors of HIV exposure at birth with a classification error of 35% using latent variable-1 (LV-1) loadings (Figure 5B). Receiver Operating Curve (ROC) analysis using LV-1 loadings showed good discrimination between iHEU and iHUU with a highly significant (p=0.004) AUC of 0.85 (Figure 5E).

**Figure 5:**
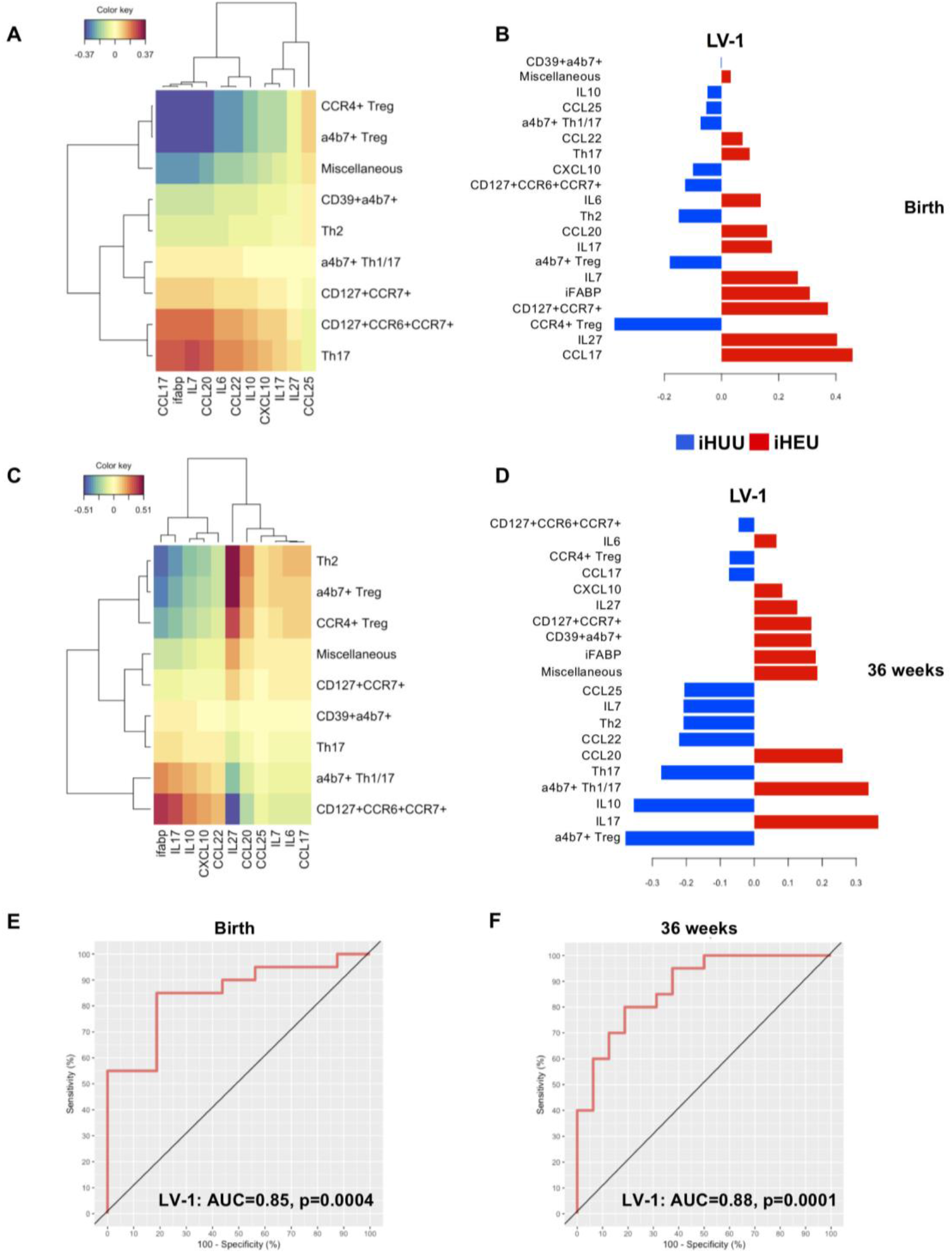
Predictive measures of HIV exposure at birth and week 36. A) Cluster image map showing RV coefficients of CD4 clusters with iFABP, chemokine and cytokine concentrations at birth. B) Sparse Partial Least Square discriminant analysis (sPLS-DA) between iHUU and iHEU using CD4 clusters and cytokine/chemokine data set at birth. C) Cluster image map showing RV coefficients of CD4 clusters with iFABP, chemokine and cytokine concentrations at 36 weeks. D) sPLS-DA between iHUU and iHEU using CD4 clusters and cytokine/chemokine data at 36 weeks. E) Receiver Operating Curve (ROC) analysis of LV-1 loadings at birth. F) ROC analysis of LV-1 loadings at week 36 of age.

At 36 weeks after birth, although no differences in chemokine and cytokine concentrations were observed (Figure 4B and S4B), there were correlations between these analytes and CD4+ T cell clusters that were similar to those found at birth (Figure 5C). The frequencies of Treg clusters (CCR4^+^ Treg and α4β7^+^ Treg) and the Th2 clusters were positively correlated with IL27 (RV=0.34, 0.47 and 0.51 respectively) and inversely correlated to iFABP (RV= −0.33, −0.42 and −0.45 respectively), whereas CD127^+^CCR6^+^CCR7^+^ and α4β7^+^ Th1/17 clusters were negatively correlated to IL27 (RV= −0.48 and −0.29 respectively) and positively correlated to iFABP (RV= 0.42 and 0.25 respectively, Figure 5C). ROC analysis at week 36 showed that the strongest predictors of HIV exposure were α4β7^+^ Treg and α4β7^+^ Th1/17 cell clusters together with IL17 and IL10, although having a higher classification error of 51% and LV-1 AUC=0.88 (Figures 5D and F).

## Discussion

This study demonstrates the intricate homeostatic balance of CD4^+^ T cell subsets during infancy that are necessary for appropriate responses to neoantigens, including pathogens, non-inherited maternal antigens, gut commensals, and autoantigens. Although serving opposite roles during an inflammatory response, Th17 and Treg cells share similar features; both are dependent on TGF-β and IL-2 for their differentiation (36), exhibit specificity towards commensal-derived antigens and are abundant in the intestine to regulate homeostasis between the immune system and commensal microbes (37). Overall, we observed a synchronous expansion of both Treg and Th17 cells early in life of healthy infants consistent with other reported studies (20, 38, 39). The greatest increase observed was within the first 7 weeks of life suggesting a period of drastic changes as the infants are exposed to diverse environmental antigens and childhood vaccines and thus requires both the regulatory and inflammatory arms of the immune system.

We further demonstrate that HIV/ART exposure disrupts the emergent Th17-Treg homeostatic balance most likely via gut epithelial damage and potential homing of Treg cells to the gut at birth. It has previously been reported that iHEU, tend to display heightened immune activation compared to iHUU (30–32), and we hypothesize from our data that this immune activation results from loss of balance between activation and tolerogenic signals. We provide evidence that the depressed Treg cells at birth in iHEU relative to iHUU are likely due to cells migrating out of circulation possibly to mitigate epithelial gut damage. Here we postulate that the highly suppressive CCR4^+^ Treg cells would be attracted to the site of inflammation via the CCL17 chemokine, a ligand for CCR4, most likely secreted by inflamed gut mucosal epithelial cells (40). This hypothesis is substantiated by high levels of CCL17 and iFABP in plasma of iHEU compared to iHUU, while the frequency of CCR4^+^ Treg cells were lower in iHEU compared to iHUU. Furthermore, Treg cells were inversely correlated to CCL17 and iFABP, and thus impaired epithelial gut cells could induce the secretion of CCL17 resulting in increased systemic CCL17 concentrations and triggering the migration of Treg cells to site of damage. Collectively, these events result in delayed peripheral ontogeny of these cells in iHEU, although by 7 weeks, iFABP and Treg cell levels are comparable between iHEU and iHUU. There is then a further progressive loss of Treg cells at 36 weeks of life, suggesting that the Th17-Treg balance may fluctuate in iHEU. Such fluctuating inflammatory-regulatory balance could in part be explained by the shifting gut microbiota profiles during weaning, feeding habits and vaccinations thus impacting gut-immune axis pre-conditioned at birth (41, 42). Studies investigating direct evidence for gastrointestinal (GI) tract damage; either though GI tract samples or endoscopy could provide further evidence for gut mucosal damage in these infants. However, these are invasive evaluations and not readily feasible in human studies of the infant.

Our dataset reveals two different Treg populations that categorize infants by HIV-exposure at birth and 36 weeks. As discussed above, events at birth are best explained by the trafficking of suppressive population of Treg cells for the reconstitution of gut epithelial damage in iHEU. Our integrated datasets could still distinguish the two groups of infants at 36 weeks of age where α4β7^+^Treg-IL10 and α4β7^+^Th1/17-IL17 were predictive of iHUU and iHEU respectively. This evidence demonstrates long-lasting effects of *in utero* HIV and/or ARV exposure on the infants’ Th17-Treg immune axis characteristic of the interplay of these immune cells in homeostatic maintenance in the gastrointestinal tract (37, 43). It is noteworthy that the observed frequency of Treg CCR4+ and α4β7^+^Th1/17 accounted for <1% of peripheral CD4^+^ cells and we speculate that these frequencies are a snapshot of CD4+ cell trafficking to mucosal sites (37).

Th17-Treg imbalance has been reported in other infant diseases associated with gut damage such as neonatal necrotizing enterocolitis, where infants presenting with disease had lower frequencies of peripheral blood Treg cells (44, 45). Non-breastfed macaque infants had elevated Th17, while mix-fed human infants exhibited activated CD4^+^ and in both studies showing that altered CD4^+^ cells was associated with changes in the gut microbiota (46, 47). Similar mechanisms could play a role in in iHEU whereby events occurring in the gut influences systemic Th17-Treg balance. Indeed, dysbiotic gut microbiota among iHEU relative to iHUU has been previously reported (48, 49). Here dysbiosis among iHEU was reported to be influenced by varying oligosaccharide composition in the breast milk of women with HIV women compared to those without HIV (48). Dybiosis in iHEU would also explain altered inflammatory condition in the gut of iHEU associated with increased epithelial permeability. This is further suggested by iHEU having higher concentrations of IL-27 albeit not significant after adjusting for multiple comparison, a regulatory cytokine involved in wound healing including intestinal barrier protection (35, 50), and thus further supported that there was some form of epithelial gut damage at birth (51, 52). Whether this altered Th17/Treg ratio is associated with loss of tolerogenic signals or an increased risk for disease among iHEU is unclear, but could in part explain the heightened risk of infections reported for iHEU compared to iHUU (12–14).

The strength of our study lies in the cohort and matching of breastfed infants born to women living with HIV to those living without HIV from the same clinic and dwelling area. However, there are several limitations. Our study was limited to a total phenotype analysis of CD4+ T cell populations based on how we collected the samples at the clinic site. Immediate fixing of whole blood only allowed us to examine surface and intracellular expression of markers associated with Th17 and Treg cells. In the absence of antigen/mitogen re-stimulation, we could not examine IL-10 or IL-17 cytokine expression to confirm Treg and Th17 cellular function. The primary focus of our study, however, was on the balance between Treg/Th17 cells and our limitation of a 15-colour panel was reflective of the deep phenotyping of such subsets. We also acknowledge that inclusion of memory differentiation markers (such as CD45RO/RA and CD27) would have allowed us to additionally understand memory ontogeny of these subsets in the first 9 months. Although this is a limitation of the present study, these changes have been described in iHEU (7–9) and are ongoing in our future studies. A further limitation of the study was the potential impact of maternal viral loads during pregnancy, birth and breastfeeding on shaping the expansion of Treg and Th17 phenotypes. We were unable to assess this due to limited sample availability. Further studies would need to investigate whether Treg cells have differing chemotactic activity between iHEU and iHUU. It is interesting to note that Jalbert et al (2019) reported higher frequencies of Treg cells in iHEU compared to iHUU (53). However, this study used cord blood and compared iHUU from a U.S cohort with iHEU from South Africa. We showed here that cord blood cellular constituents are distinct from birth peripheral blood and most likely does not reflect events in the newborn infant blood, agreeing with others (22). It is also very important to have stringent measures of Treg cells, where minimal markers would be CD4^+^ CD25^high^ Foxp3^+^ CD127^-^ (54). Our deep phenotyping panel used traditional markers for both Treg and Th17 cells, but also markers associated with highly suppressive Treg cells. For example, TIGIT and CD39 would be expressed on highly suppressive Treg cells (55), where CD39 expression on induced Treg cells, being an ectonucleatidase enzyme involved in the conversion of extracellular ATP into AMP, has been shown to alleviate IL-17- and IFNg-driven inflammatory responses (56).

The postnatal period is a critical window for priming and the maturation of the infants’ immune system and its interaction with commensal microbes to facilitate homeostasis (7, 8). Perturbations occurring during this “window of opportunity” alter ontogeny of immune development and likely associate with long-term immune-related diseases (8, 22). The association between markers of gut damage and *in utero* HIV/ARV exposure on the Th17:Treg balance at birth has allowed us to hypothesize that a nexus exists between immune ontogeny and the gut.

## Acknowledgments

The authors would like to thank all the mothers and infants who volunteered to participate in this study and the dedicated work of the INFANT team.

## Author contributions

The study was conceived and designed by SD, CAB, HBJ and CMG. SD, MSSS and AK designed flow cytometry and cell sorting experiments. BA and HBJ were responsible for participant enrolment, sample collection and processing. Data generation and acquisition was performed by SD and analysed by SD, KL, and SPH. SD drafted the original manuscript that was reviewed and edited by all authors.

## References

1. Liu, L., S. Oza, D. Hogan, Y. Chu, J. Perin, J. Zhu, J. E. Lawn, S. Cousens, C. Mathers, and R. E. Black. 2016. Global, regional, and national causes of under-5 mortality in 2000-15: an updated systematic analysis with implications for the Sustainable Development Goals. Lancet 388: 3027–3035.

2. Dowling, D. J., and O. Levy. 2014. Ontogeny of early life immunity. Trends Immunol. 35: 299–310.

3. Ivarsson, M. A., L. Loh, N. Marquardt, E. Kekäläinen, L. Berglin, N. K. Björkström, M. Westgren, D. F. Nixon, and J. Michaëlsson. 2013. Differentiation and functional regulation of human fetal NK cells. J. Clin. Invest. 123: 3889–3901.

4. Kraft, J. D., J. Horzempa, C. Davis, J. Y. Jung, M. M. O. Peña, and C. M. Robinson. 2013. Neonatal macrophages express elevated levels of interleukin-27 that oppose immune responses. Immunology 139: 484–493.

5. Mold, J. E., S. Vankatasubrahmanyam, T. D. Burt, J. Michaelsson, J. M. Rivera, S. A. Galkina, K. Weinberg, C. A. Stoddart, and J. M. McCune. 2010. Fetal and Adult Hematopoietic Stem Cells Give Rise to Distinct T Cell Lineages in Humans. Science (80.). 330: 1695–9.

6. Kollmann, T. R., O. Levy, R. R. Montgomery, and S. Goriely. 2012. Innate Immune Function by Toll-like Receptors: Distinct Responses in Newborns and the Elderly. Immunity 37: 771–783.

7. Torow, N., and M. W. Hornef. 2017. The Neonatal Window of Opportunity: Setting the Stage for Life-Long Host-Microbial Interaction and Immune Homeostasis. J. Immunol. 198: 557–563.

8. Gensollen, T., S. S. Iyer, D. L. Kasper, and R. S. Blumberg. 2016. How colonization by microbiota in early life shapes the immune system. Science (80-.). 352: 539–544.

9. Noack, M., and P. Miossec. 2014. Th17 and regulatory T cell balance in autoimmune and inflammatory diseases. Autoimmun. Rev. 13: 668–677.

10. Slogrove, A. L., K. M. Powis, L. F. Johnson, J. Stover, and M. Mahy. 2020. Estimates of the global population of children who are HIV-exposed and uninfected, 2000-18: a modelling study. Lancet Glob. Heal. 8: e67–e75.

11. Brennan, A. T., R. Bonawitz, C. J. Gill, D. M. Thea, M. Kleinman, J. Useem, L. Garrison, R. Ceccarelli, C. Udokwu, L. Long, and M. P. Fox. 2016. A meta-analysis assessing all-cause mortality in HIVexposed uninfected compared with HIV-unexposed uninfected infants and children. Aids 30: 2351–2360.

12. Cohen, C., J. Moyes, S. Tempia, M. Groome, S. Walaza, M. Pretorius, F. Naby, O. Mekgoe, K. Kahn, A. von Gotterberg, N. Wolter, A. Cohenm, C. von Mollendorf, M. Venter, and S. A. Madhi. 2016. Epidemiology of Acute Lower Respiratory Tract Infection in HIV-Exposed Uninfected Infants. Pediatrics 137: e20153272.

13. Slogrove, A. L., T. Goetghebuer, M. F. Cotton, J. Singer, and J. Bettinger. 2016. Pattern of Infectious Morbidity in HIV-Exposed Uninfected Infants and Children. Front. Immunol. 7: 1–8.

14. Brennan, A. T., R. Bonawitz, C. J. Gill, D. M. Thea, M. Kleinman, L. Long, C. McCallum, and M. P. Fox. 2019. A meta-analysis assessing diarrhea and pneumonia in HIV-exposed uninfected compared to HIV-unexposed uninfected infants and children. JAIDS J. Acquir. Immune Defic. Syndr. 82: 1–8.

15. Tchakoute, C. T., K. L. Sainani, S. Osawe, P. Datong, A. Kiravu, K. L. Rosenthal, C. M. Gray, D. W. Cameron, A. Abimiku, and H. B. Jaspan. 2018. Breastfeeding mitigates the effects of maternal HIV on infant infectious morbidity in the Option B+ era. AIDS 32: 2383–2391.

16. World Health Organisation. 2016. Consolidated guidelines on the use of antiretroviral drugs for treating and preventing HIV infection: recommendations for a public health approach. World Heal. Organ.

17. Seiler, C., A.-M. Ferreira, L. M. Kronstad, L. J. Simpson, M. Le Gars, E. Vendrame, C. A. Blish, and S. Holmes. 2021. CytoGLMM: conditional differential analysis for flow and mass cytometry experiments. BMC Bioinformatics 22: 1–14.

18. Nowicka, M., C. Krieg, H. Crowell, L. M. Weber, F. J. Hartmann, S. Guglietta, B. Becher, M. P. Levesque, and M. D. Robinson. 2017. CyTOF workflow: Differential discovery in high-throughput high-dimensional cytometry datasets. F1000Research 6.

19. Rohart, F., B. Gautier, A. Singh, and K. A. Lê Cao. 2017. mixOmics: An R package for ‘omics feature selection and multiple data integration. PLoS Comput. Biol. 13: 1–19.

20. Collier, F. M., M. K. Tang, D. Martino, R. Saffery, J. Carlin, K. Jachno, S. Ranganathan, D. Burgner, K. J. Allen, P. Vuillermin, and A.-L. Ponsonby. 2015. The ontogeny of naive and regulatory CD4+ T-cell subsets duing the first postnatal year: a cohort study. Clin. Transl. Immunol. 4: e34.

21. Dirix, V., F. Vermeulen, and F. Mascart. 2013. Maturation of CD4+ regulatory T lymphocytes and of cytokine secretions in infants born prematurely. J. Clin. Immunol. 33: 1126–1133.

22. Olin, A., E. Henckel, Y. Chen, T. Lakshmikanth, C. Pou, J. Mikes, A. Gustafsson, A. K. Bernhardsson, C. Zhang, K. Bohlin, and P. Brodin. 2018. Stereotypic Immune System Development in Newborn Children. Cell 174: 1277–1292.e14.

23. Sehrawat, S., and B. T. Rouse. 2017. Interplay of regulatory T cell and Th17 cells during infectious diseases in humans and animals. Front. Immunol. 8: 1–14.

24. Gagliani, N., M. C. A. Vesely, A. Iseppon, L. Brockmann, H. Xu, N. W. Palm, M. R. de Zoete, P. Licona-Limón, R. S. Paiva, T. Ching, C. Weaver, X. Zi, X. Pan, R. Fan, L. X. Garmire, M. J. Cotton, Y. Drier, B. Bernstein, J. Geginat, B. Stockinger, E. Esplugues, S. Huber, and R. A. Flavell. 2015. Th17 cells transdifferentiate into regulatory T cells during resolution of inflammation. Nature 523: 221–5.

25. Bystrom, J., F. I. L. Clanchy, T. E. Taher, M. Al-Bogami, V. H. Ong, D. J. Abraham, R. O. Williams, and R. A. Mageed. 2019. Functional and phenotypic heterogeneity of Th17 cells in health and disease. Eur. J. Clin. Invest. 49: 1–13.

26. Matos, T. R., M. Hirakawa, A. C. Alho, L. Neleman, L. Graca, and J. Ritz. 2021. Maturation and Phenotypic Heterogeneity of Human CD4+ Regulatory T Cells From Birth to Adulthood and After Allogeneic Stem Cell Transplantation. Front. Immunol. 11: 1–10.

27. Borsellino, G., M. Kleinewietfeld, D. Di Mitri, A. Sternjak, A. Diamantini, R. Giometto, S. Hopner, C. Diego, G. Bernardi, M. L. Dell’Acwua, P. M. Rossini, L. Battistini, O. Rotzschke, and K. Falk. 2007. Expression of ectonucleotidase CD39 by Foxp3+ Treg cells: hydrolysis of extracellular ATP and immune suppression. Blood 110: 1225–1232.

28. Joller, N., E. Lozano, P. R. Burkett, B. Patel, S. Xiao, C. Zhu, J. Xia, T. G. Tan, E. Sefik, V. Yajnik, A. H. Sharpe, J. Francisco, D. Mathis, C. Benoist, D. A. Hafler, and V. K. Kuchroo. 2014. Treg cells expressing the co-inhibitory molecule TIGIT selectively inhibity pro-inflammatory Th1 and Th17 cell responses. Immunity 40: 569–581.

29. Reikie, B. A., R. C. M. Adams, A. Leligdowicz, K. Ho, S. Naidoo, C. E. Rusk, C. De Beer, and T. R. Kollmann. 2014. Altered innante immune development in HIV-exposed uninfected infants. J. Acquir. Immune Defic. Syndriome 66: 245–255.

30. Rich, K. C., J. N. Siegel, C. Jennings, R. J. Rydman, and a L. Landay. 1997. Function and phenotype of immature CD4+ lymphocytes in healthy infants and early lymphocyte activation in uninfected infants of human immunodeficiency virus-infected mothers. Clin. Diagn. Lab. Immunol. 4: 358–361.

31. Clerici, M., M. Saresella, F. Colombo, S. Fossati, N. Sala, D. Bricalli, M. L. Villa, P. Ferrante, L. Dally, A. Vigano, and W. Dc. 2000. T-lymphocyte maturation abnormalities in uninfected newborns and children with vertical exposure to HIV T-lymphocyte maturation abnormalities in uninfected newborns and children with vertical exposure to HIV. Bloid 96: 3866–3871.

32. Rainwater-Lovett, K., N. Hc, M. Mubiana-Mbewe, B. Moore C, M. Jb, and M. Wj. 2014. Changes in Cellular Immune Activation and Memory T Cell Subsets in HIV-Infected Zambian Children Receiving HAART. J. Acquir. Immune Defic. Syndr. 67: 455–462.

33. Prendergast, A. J., B. Chasekwa, S. Rukobo, M. Govha, K. Mutasa, R. Ntozini, and J. H. Humphrey. 2017. Intestinal Damage and Inflammatory Biomarkers in Human Immunodeficiency Virus (HIV)-Exposed and HIV-Infected Zimbabwean Infants. J. Infect. Dis. 216: 651–661.

34. Wells, J. M., R. J. Brummer, M. Derrien, T. T. MacDonald, F. Troost, P. D. Cani, V. Theodorou, J. Dekker, A. Méheust, W. M. De Vos, A. Mercenier, A. Nauta, and C. L. Garcia-Rodenas. 2017. Homeostasis of the gut barrier and potential biomarkers. Am. J. Physiol. - Gastrointest. Liver Physiol. 312: G171–G193.

35. Diegelmann, J., T. Olszak, B. Göke, R. S. Blumberg, and S. Brand. 2012. A novel role for interleukin-27 (IL-27) as mediator of intestinal epithelial barrier protection mediated via differential signal transducer and activator of transcription (STAT) protein signaling and induction of antibacterial and anti-inflammatory protei. J. Biol. Chem. 287: 286–298.

36. Weaver, C. T., L. E. Harrington, P. R. Mangan, M. Gavrieli, and K. M. Murphy. 2006. Th17: An Effector CD4 T Cell Lineage with Regulatory T Cell Ties. Immunity 24: 677–688.

37. Shen, X., J. Du, W. Guan, and Y. Zhao. 2014. The balance of intestinal Foxp3+ regulatory T cells and Th17 cells and its biological significance. Expert Rev. Clin. Immunol. 10: 353–362.

38. Black, A., S. Bhaumik, R. L. Kirkman, C. T. Weaver, and D. A. Randolph. 2012. Developmental regulation of Th17-cell capacity in human neonates. Eur. J. Immunol. 42: 311–319.

39. Hayakawa, S., N. Ohno, S. Okada, and M. Kobayashi. 2017. Significant augmentation of regulatory T cell numbers occurs during the early neonatal period. Clin. Exp. Immunol. 190: 268–279.

40. Heiseke, A. F., A. C. Faul, H. Lehr, I. Förster, R. M. Schmid, A. B. Krug, and W. Reindl. 2012. CCL17 promotes intestinal inflammation in mice and counteracts regulatory T cellmediated protection from colitis. Gastroenterology 142: 335–345.

41. Bäckhed, F., J. Roswall, Y. Peng, Q. Feng, H. Jia, P. Kovatcheva-Datchary, Y. Li, Y. Xia, H. Xie, H. Zhong, M. T. Khan, J. Zhang, J. Li, L. Xiao, J. Al-Aama, D. Zhang, Y. S. Lee, D. Kotowska, C. Colding, V. Tremaroli, Y. Yin, S. Bergman, X. Xu, L. Madsen, K. Kristiansen, J. Dahlgren, and W. Jun. 2015. Dynamics and stabilization of the human gut microbiome during the first year of life. Cell Host Microbe 17: 690–703.

42. Al Nabhani, Z., S. Dulauroy, R. Marques, C. Cousu, S. Al Bounny, F. Déjardin, T. Sparwasser, M. Bérard, N. Cerf-Bensussan, and G. Eberl. 2019. A Weaning Reaction to Microbiota Is Required for Resistance to Immunopathologies in the Adult. Immunity 50: 1276–1288.e5.

43. Omenetti, S., and T. T. Pizarro. 2015. The Treg/Th17 axis: A dynamic balance regulated by the gut microbiome. Front. Immunol. 6: 1–8.

44. Pang, Y., X. Du, X. Xu, M. Wang, and Z. Li. 2018. Impairment of regulatory T cells in patients with neonatal necrotizing enterocolitis. Int. Immunopharmacol. 63: 19–25.

45. Pang, Y., X. Du, X. Xu, M. Wang, and Z. Li. 2018. Monocyte activation and inflammation can exacerbate Treg/Th17 imbalance in infants with neonatal necrotizing enterocolitis. Int. Immunopharmacol. 59: 354–360.

46. Wood, L. F., B. P. Brown, K. Lennard, U. Karaoz, E. Havyarimana, J. A. S. Passmore, A. C. Hesseling, P. T. Edlefsen, L. Kuhn, N. Mulder, E. L. Brodie, D. L. Sodora, and H. B. Jaspan. 2018. Feeding-Related Gut Microbial Composition Associates with Peripheral T-Cell Activation and Mucosal Gene Expression in African Infants. Clin. Infect. Dis. 67: 1237–1246.

47. Ardeshir, A., N. R. Narayan, G. Méndez-Lagares, D. Lu, M. Rauch, Y. Huang, K. K. A. Van Rompay, S. V. Lynch, and D. J. Hartigan-O’Connor. 2014. Breast-fed and bottle-fed infant rhesus macaques develop distinct gut microbiotas and immune systems. Sci. Transl. Med. 6.

48. Bender, J. M., F. Li, S. Martelly, E. Byrt, V. Rouzier, M. Leo, N. Tobin, P. S. Pannaraj, H. Adisetiyo, A. Rollie, C. Santiskulvong, S. Wang, C. Autran, L. Bode, D. Fitzgerald, L. Kuhn, and G. M. Aldrovandi. 2016. Maternal HIV infection influences the microbiome of HIV-uninfected infants. Sci. Transl. Med. 8: 349ra100.

49. Machiavelli, A., R. T. D. Duarte, M. M. d. S. Pires, C. R. Zárate-Bladés, and A. R. Pinto. 2019. The impact of in utero HIV exposure on gut microbiota, inflammation, and microbial translocation. Gut Microbes 10: 599–614.

50. Yang, B., J. Suwanpradid, R. Sanchez-Lagunes, H. W. Choi, P. Hoang, D. Wang, S. N. Abraham, and A. S. MacLeod. 2017. IL-27 Facilitates Skin Wound Healing through Induction of Epidermal Proliferation and Host Defense. J. Invest. Dermatol. 137: 1166—1175.

51. Papasavvas, E., L. Azzoni, A. Foulkes, A. Violari, M. F. Cotton, M. Pistilli, G. Reynolds, X. Yin, D. K. Glencross, W. S. Stevens, J. A. McIntyre, and L. J. Montaner. 2011. Increased microbial translocation in ≤180 days old perinatally human immunodeficiency virus-positive infants as compared with human immunodeficiency virus-exposed uninfected infants of similar age. Pediatr. Infect. Dis. J. 30: 877–882.

52. Pilakka-Kanthikeel, S., A. Kris, A. Selvaraj, S. Swaminathan, and S. Pahwa. 2014. Immune activation is associated with increased gut microbial translocation in treatment-naive, HIV-infected children in a resource-limited setting. J. Acquir. Immune Defic. Syndr. 66: 16–24.

53. Jalbert, E., K. M. Williamson, M. E. Kroehl, M. J. Johnson, C. Cutland, S. A. Madhi, M. C. Nunes, and A. Weinberg. 2019. HIV-exposed uninfected infants have increased regulatory T cells that correlate with decreased T cell function. Front. Immunol. 10: 1–9.

54. Santegoets, S. J. A. M., E. M. Dijkgraaf, A. Battaglia, P. Beckhove, C. M. Britten, A. Gallimore, A. Godkin, C. Gouttefangeas, T. D. de Gruijl, H. J. P. M. Koenen, A. Scheffold, E. M. Shevach, J. Staats, K. Taskén, T. L. Whiteside, J. R. Kroep, M. J. P. Welters, and S. H. van der Burg. 2015. Monitoring regulatory T cells in clinical samples: consensus on an essential marker set and gating strategy for regulatory T cell analysis by flow cytometry. Cancer Immunol. Immunother. 64: 1271–1286.

55. Salvany-Celades, M., A. van der Zwan, M. Benner, V. Setrajcic-Dragos, H. A. Bougleux Gomes, V. Iyer, E. R. Norwitz, J. L. Strominger, and T. Tilburgs. 2019. Three Types of Functional Regulatory T Cells Control T Cell Responses at the Human Maternal-Fetal Interface. Cell Rep. 27: 2537–2547.e5.

56. Su, W., X. Chen, W. Zhu, J. Yu, W. Li, Y. Li, Z. Li, N. Olsen, D. Liang, and S. G. Zheng. 2019. The cAMP–Adenosine Feedback Loop Maintains the Suppressive Function of Regulatory T Cells. J. Immunol. 203: 1436–1446.

