## Supplementary information for "Stereotypic expansion of T_regulatory_ and Th_17_ cells during infancy is disrupted by HIV exposure and gut epithelial damage"

**Table Legends**

**Table S1** Demographic characteristic of HIV infected-exposed and HIV uninfected-unexposed mother-infant pairs.

**Table S2** Flow cytometry antibody panel used to surface and intracellular stain infant whole blood to characterize T cells.

**Figure Legends**

**Figure S1: Flow cytometry gating strategy to identify T regulatory (Treg) and Th17 cells**. Fixed whole blood cells stained with the 15-color panel showing the singlet gate (to exclude doublets), lymphocyte gate and a CD3+/live-dead (VIVID) gate to identify viable CD4 and CD8 subsets. CD4+ T cells gates was used to define Th17 cells as CCR6+CCR4+CD161+ cells. Also shown are the gated CD4 cells, co-expressing CD25 dim (green gate) and bright (red gate) in the absence or presence (purple) CD127 expression. The overlay of these gates onto univariate plots of FoxP3, CD39 and TIGIT are shown.

**Figure S2: Changes in Th17 and Treg CD4$^{+}$ T cell clusters using SOM analysis in HIV-exposed uninfected (iHEU) and HIV-unexposed uninfected (iHUU) infants.** A) Comparing frequencies of Th17 and Treg CD4$^{+}$ clusters between iHEU and iHUU at birth, 7, 15 and 36 weeks. B) Differences in the frequencies of Th17 and Treg CD4$^{+}$ clusters in iHEU between mothers who were initiated on cART prior pregnancy (stable: iHEU-s) and those initiated cART during pregnancy (initiating: iHEU-i). This was used as a proxy for potential HIV exposure. Statistical comparisons were made using the Mann-Whitney *U* test.

**Figure S3: Changes in Th17 and Treg CD4$^+$ T cell manually gated phenotypes in HIV-exposed uninfected (iHEU) and HIV-unexposed uninfected (iHUU) infants.** A and B) Comparing frequencies of Th17 and Treg CD4$^+$ clusters between iHEU and iHUU at birth, 7, 15 and 36 weeks. C and D) Differences in the frequencies of Th17 and Treg CD4$^+$ manually gated cell frequencies in iHEU between mothers who were initiated on cART prior pregnancy (stable: iHEU-s) and those initiated cART during pregnancy (initiating: iHEU-i). This was used as a proxy for potential HIV exposure. Statistical comparisons were made using the Mann-Whitney *U* test.

**Figure S4**: $Log_2$ Th17/Treg ratio in iHEU and iHUU at birth, 7, 15 and 36 weeks of age. Friedman's test was used to show changes within a group and * $p<0.05$ Mann-Whitney *U* test used to compare between groups.

**Figure S5: Plasma analytes measured in HIV-exposed uninfected infants (iHEU) and HIV-unexposed uninfected infants (iHUU).** A) Comparisons of iFABP concentrations between iHEU and iHUU measured at 7 and 15 weeks. B). Comparison of plasma iFABP concentrations in iHEU stratified by timing of maternal combined antiretroviral treatment (cART) during pregnancy and mothers initiating cART prior to pregnancy (stable: iHEU-s). C) Comparisons of chemokine and cytokine concentrations between iHEU and iHUU measured at birth and 36 weeks of age. D) Chemokine and cytokine concentrations in iHEU stratified by timing of maternal cART (iHEU-s vs iHEU-i). Statistical comparisons were made using the Mann-Whitney *U* test.

Table S1 Demographic characteristic of HIV-uninfected and HIV-infected mothers and their respective HIV uninfected-unexposed infants (iHUU) and HIV-exposed uninfected infants (iHEU).

| **Demographics** | **N=16** | **N=20** | | **P value** |
|---|---|---|---|---|
| **Mother** | HIV-uninfected | HIV-infected | | |
| Median maternal age at delivery, years (range) | 26 (19-36) | 27 (19-39) | | 0.49 |
| Median gestational age, weeks (range) | 39 (37-41) | 39 (37-41) | | 0.89 |
| Median maternal CD4 count cells/µL (IQR) | - | 429.5 (344.5-516.50) | | |
| Initiating cART during pregnancy, n (%) | | 10 (50) | | |
| | | cART-i* | cART-s* | |
| Median duration of cART during pregnancy, weeks (IQR) | | 24.9 (19.8-28.6) | 39.0 (38.0-39.8) | |
| Median maternal CD4 count cells/µL (IQR) | | 388.5 (330.8-501.8) | 474.5 (389.0-508.0) | 0.49 |
| Infant | iHUU | iHEU | | |
| Median birth weight, kg (IQR) | 3.0 (2.9-3.3) | 3.3 (2.9-3.5) | | 0.31 |
| Gender, female (%) | 9 (56.3) | 11 (55.0%) | | 0.94 |
| Exclusively breastfeeding | | | | |
| 7 weeks, n (%) | 9 (56.3) | 10 (50.0) | | |
| 15 weeks, n (%) | 6 (37.5) | 6 (30.0) | | |
| 36 weeks, n (%) | 0 (0.0) | 0 (0.0) | | |

* cART-i = initiated combined antiretroviral treatment (cART) during pregnancy, ART-s = commenced cART prior pregnancy.

Table S2 Flow cytometry antibody panel used to surface and intracellular stain infant whole blood to characterize T cells.

Table S2 Flow cytometry antibody panel used to surface and intracellular stain infant whole blood to characterize T cells.

| Antibodies | Clone | Supplier | Characteristics | Purpose |
|---|---|---|---|---|
| CD3 Apc-Cy7 | UCHTI | Biolegend | | T cell markers |
| CD4 AlexaFluor 700 | SK3 | Biolegend | | |
| CD8 BV570 | RPA-T8 | Biolegend | | |
| CD127 PE-CF594 | hIL-7R-M21 | BD | IL-7Rα | T regulatory cell markers |
| CD25 PE-Cy7 | 2A3 | BD | IL-2Rα | |
| Foxp3 AlexaFluor 647 | 2594/C7 | BD | Treg transcription factor | |
| CD39 BV786 | Tu66 | BD | ectonucleatidase enzyme | |
| TIGIT FITC | MBSA43 | eBioscience | Co-inhibitory molecule | Th17 cell markers |
| CCR6 BV711 | G034E3 | eBioscience | CCL20 | |
| CCR4 BV605 | L291H4 | Biolegend | CCL17/CCL22 | |
| CD161 PE-Cy5 | DX12 | BD | | |
| CCR7 PerCP-Cy5.5 | G043H7 | Biolegend | | Lymph node tropic marker |
| α4β7 PE | A4B7R1 | NIH AIDS reagents | | Gut-tropic marker |

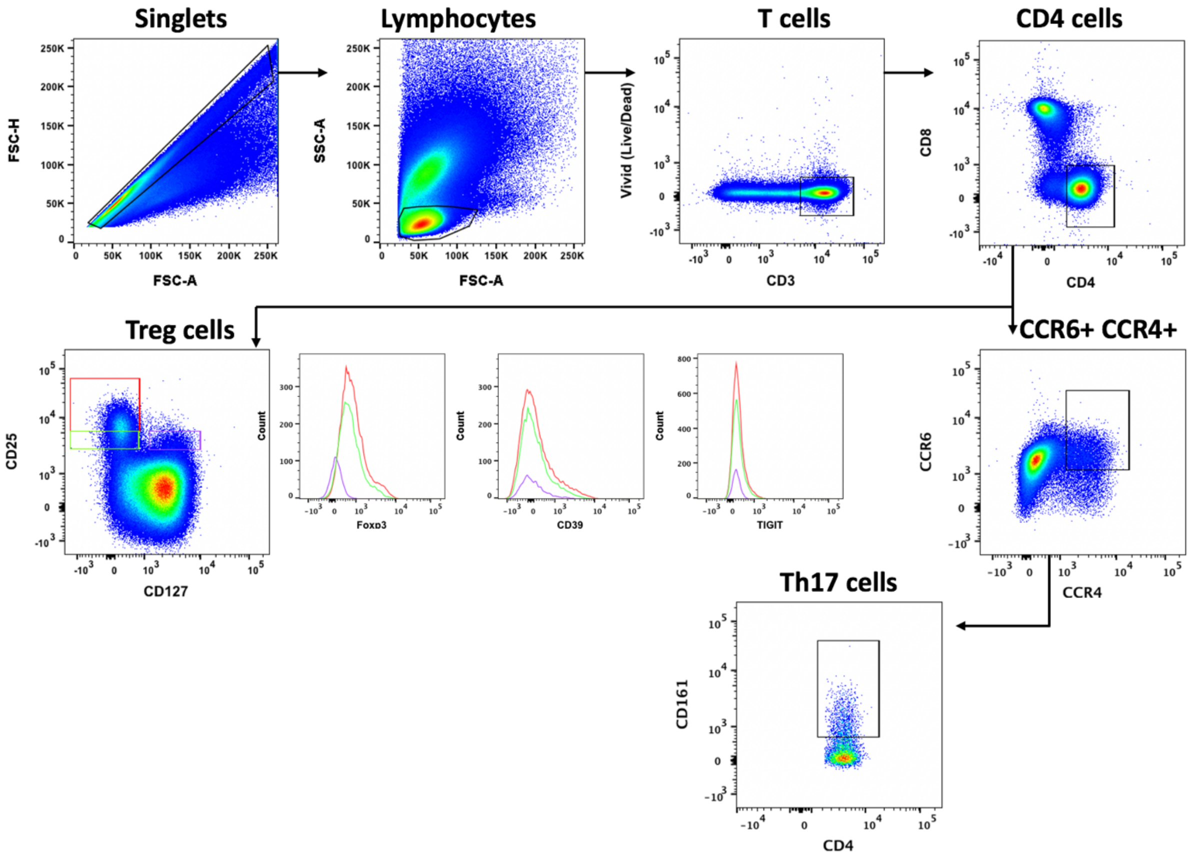


**Figure S1: Flow cytometry gating strategy to identify T regulatory (Treg) and Th17 cells**. Fixed whole blood cells stained with the 15-color panel showing the singlet gate (to exclude doublets), lymphocyte gate and a CD3+/live-dead (VIVID) gate to identify viable CD4 and CD8 subsets. CD4+ T cells gates was used to define Th17 cells as CCR6+CCR4+CD161+ cells. Also shown are the gated CD4 cells, co-expressing CD25 dim (green gate) and bright (red gate) in the absence or presence (purple) CD127 expression. The overlay of these gates onto univariate plots of FoxP3, CD39 and TIGIT are shown.

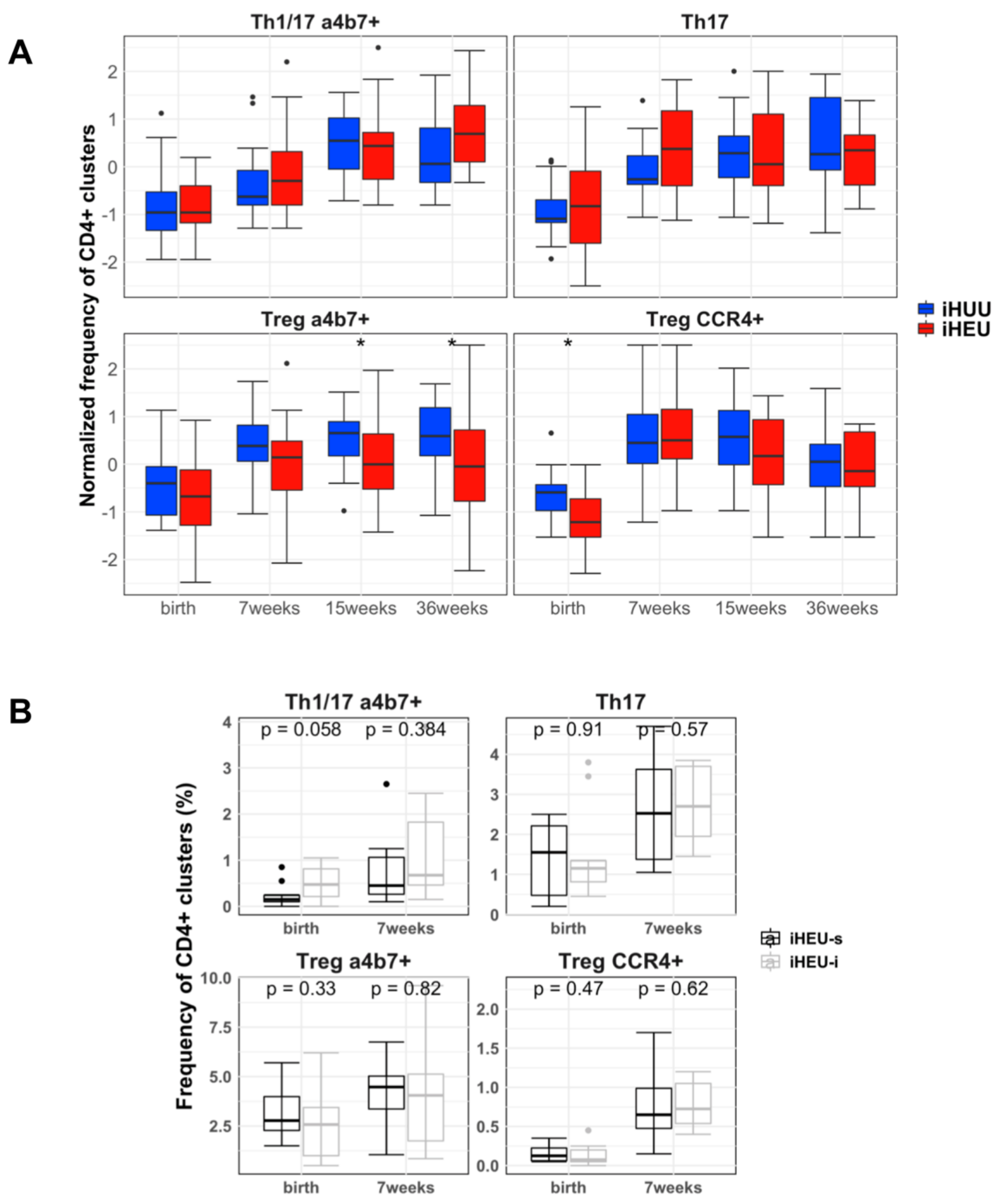


**Figure S2: Changes in Th17 and Treg CD4$^{+}$ T cell clusters using SOM analysis in HIV-exposed uninfected (iHEU) and HIV-unexposed uninfected (iHUU) infants.** A) Comparing frequencies of Th17 and Treg CD4$^{+}$ clusters between iHEU and iHUU at birth, 7, 15 and 36 weeks. B) Differences in the frequencies of Th17 and Treg CD4$^{+}$ clusters in iHEU between mothers who were initiated on cART prior pregnancy (stable: iHEU-s) and those initiated cART during pregnancy (initiating: iHEU-i). This was used as a proxy for potential HIV exposure. Statistical comparisons were made using the Mann-Whitney *U* test.

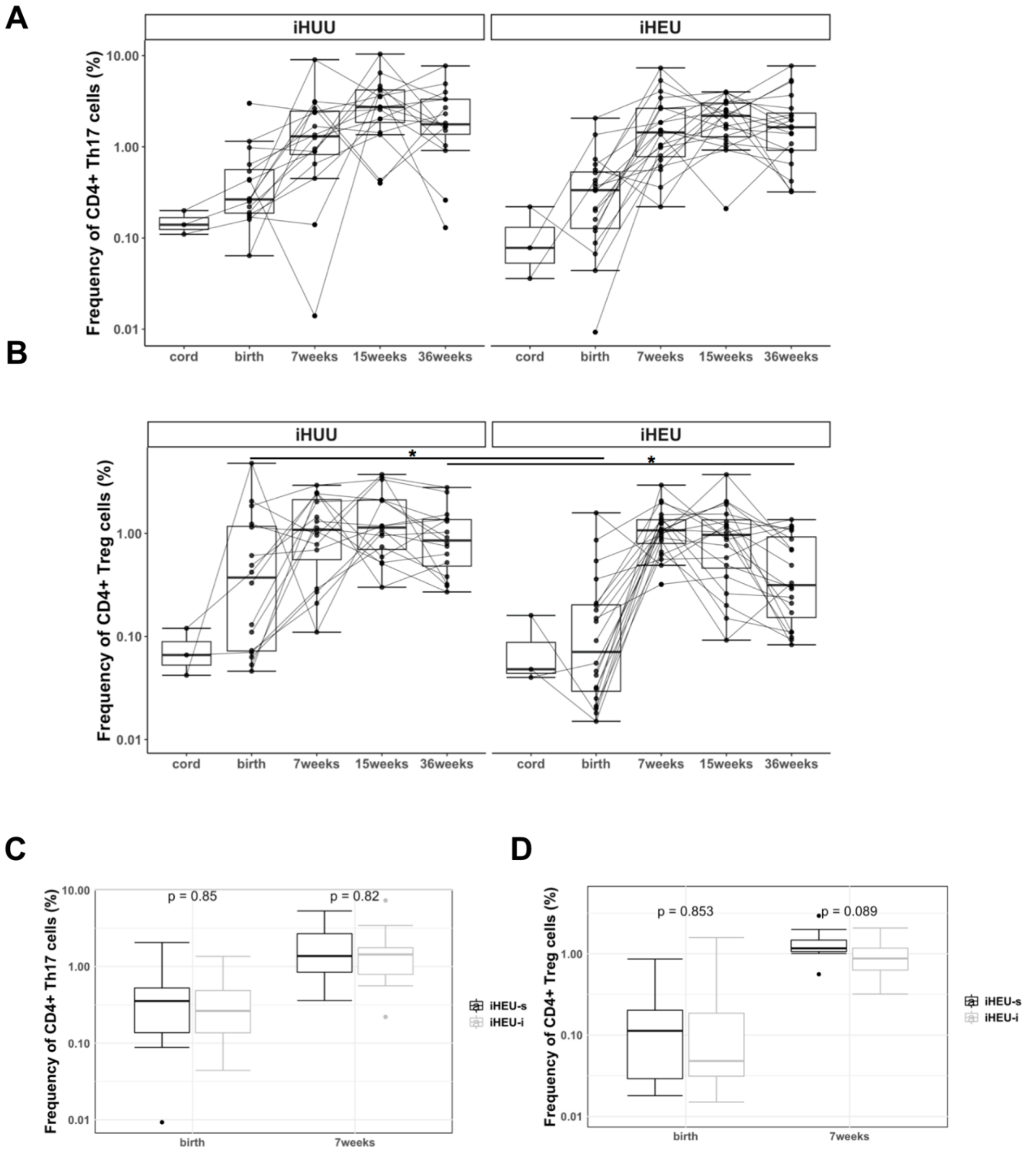


**Figure S3: Changes in Th17 and Treg CD4$^{+}$ T cell manually gated phenotypes in HIV-exposed uninfected (iHEU) and HIV-unexposed uninfected (iHUU) infants.** A and B) Comparing frequencies of Th17 and Treg CD4$^{+}$ clusters between iHEU and iHUU at birth, 7, 15 and 36 weeks. C and D) Differences in the frequencies of Th17 and Treg CD4$^{+}$ manually gated cell frequencies in iHEU between mothers who were initiated on cART prior pregnancy (stable: iHEU-s) and those initiated cART during pregnancy (initiating: iHEU-i). This was used as a proxy for potential HIV exposure. Statistical comparisons were made using the Mann-Whitney *U* test.

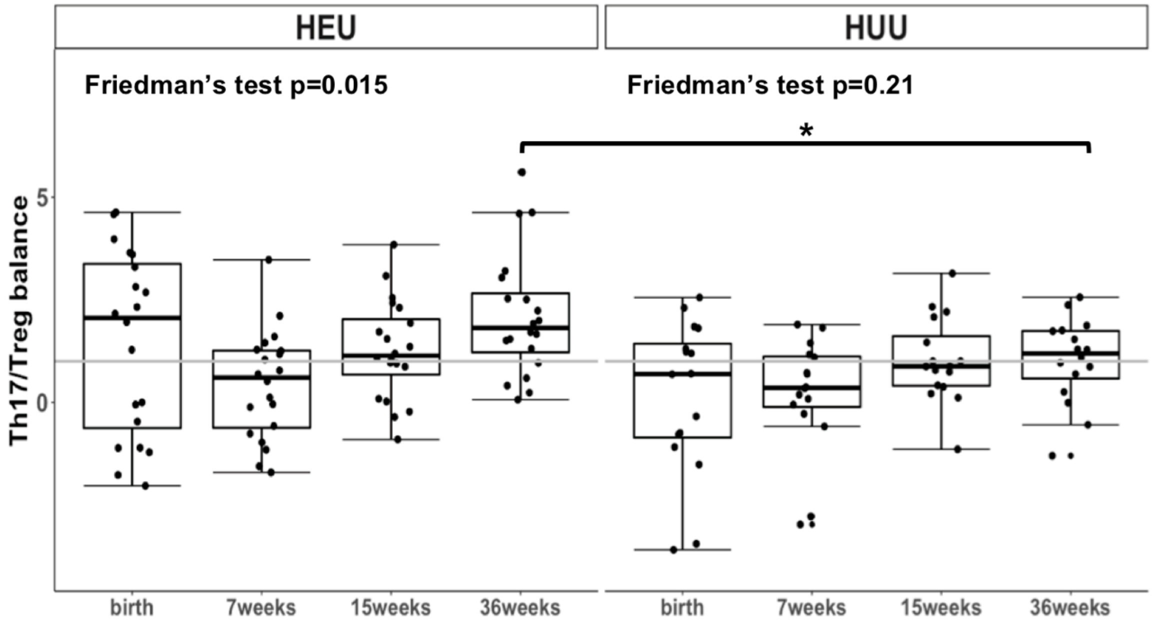


**Figure S4**: Log$_2$ Th17/Treg ratio in iHEU and iHUU at birth, 7, 15 and 36 weeks of age. Friedman's test was used to show changes within a group and * p<0.05 Mann-Whitney *U* test used to compare between groups.

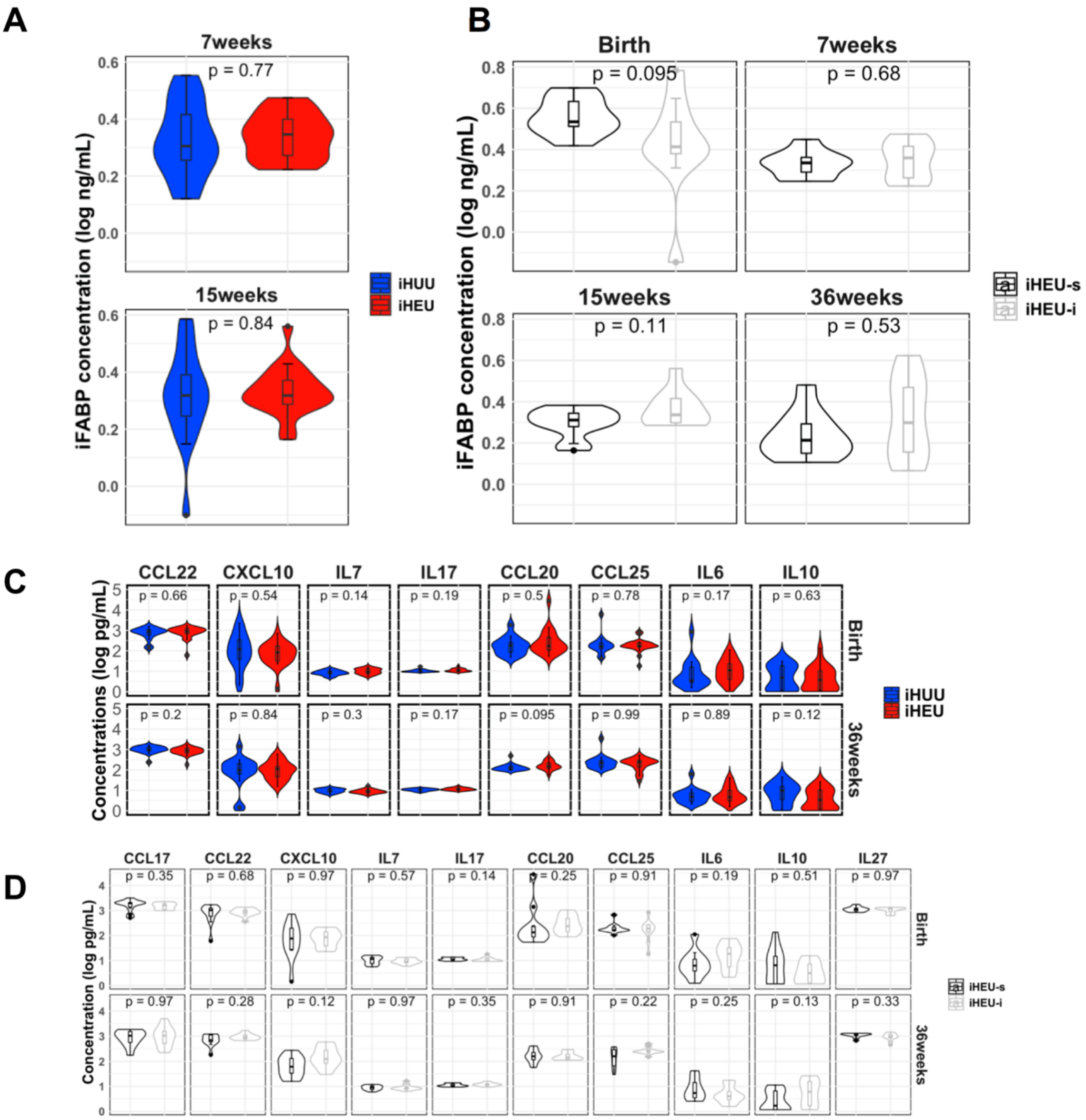


**Figure S5: Plasma analytes measured in HIV-exposed uninfected infants (iHEU) and HIV-unexposed uninfected infants (iHUU).** A) Comparisons of iFABP concentrations between iHEU and iHUU measured at 7 and 15 weeks. B). Comparison of plasma iFABP concentrations in iHEU stratified by timing of maternal combined antiretroviral treatment (cART) during pregnancy and mothers initiating cART prior to pregnancy (stable: iHEU-s). C) Comparisons of chemokine and cytokine concentrations between iHEU and iHUU measured at birth and 36 weeks of age. D) Chemokine and cytokine concentrations in iHEU stratified by timing of maternal cART (iHEU-s vs iHEU-i). Statistical comparisons were made using the Mann-Whitney *U* test.
